# Analysis of the H-Ras mobility pattern *in vivo* shows cellular heterogeneity inside epidermal tissue

**DOI:** 10.1101/2021.06.21.449125

**Authors:** Radoslaw J. Gora, Babette de Jong, Patrick van Hage, Mary A. Rhiemus, Fjodor van Steenis, John van Noort, Thomas Schmidt, Marcel J.M. Schaaf

## Abstract

Over the last two decades, developments in single-molecule microscopy (SMM) have enabled imaging and tracking of individual, fluorescently labelled proteins in biological systems, and most of these studies have focused on the analysis of protein mobility patterns inside cultured cells. In the present study, SMM was applied *in vivo*, using the zebrafish embryo model. We studied the protein dynamics of the membrane protein H-Ras, mutants of this protein, and its membrane-anchoring domain, C10H-Ras, in epidermal cells of living two-day-old embryos, using a total internal reflection fluorescence microscopy (TIRFM) setup. For all proteins studied, our results consistently confirm the presence of a fast- and a slow-diffusing subpopulations of molecules, which both confine to microdomains within the plasma membrane. Although the mobility patterns of H-Ras, mutants of this proteins and its membrane-anchoring domain were remarkably similar, the constitutively active H-Ras mutant, H-Ras^V12^, exhibited significantly higher diffusion rates than the wild-type H-Ras and its inactive mutant, H-Ras^N17^. Ultimately, we characterized variability in our data obtained using the zebrafish embryo model and demonstrated that differences between cells within the same embryo are the largest source of variation in our data. Our findings are in line with a model in which the cellular architecture of individual cells within a tissue determine the mobility of H-Ras proteins anchored in the plasma membrane cytoplasmic leaflet. Thereby, our results underline the growing importance of performing SMM imaging *in vivo* in order to better understand factors influencing the protein dynamics in an intact living organism.

**SUMMARY STATEMENT:** By application of single-molecule microscopy to living zebrafish embryos, factors altering the *in vivo* dynamics of H-Ras proteins in epidermal cells were analyzed.

## INTRODUCTION

Plasma membranes mostly consist of proteins and lipids that move laterally within the fluidic membrane plane and interact with each other in both random and organized manners (Singer and Nicolson, 1972). Until now, it has remained unclear what the origins are of many of such interactions and how the organization and mobility of all membrane constituents are governed (Gheber, 2018). Furthermore, not much is known about the role of structural complexes, such as the subjacent actomyosin cortex, in the dynamics of proteins and lipids in the plasma membrane. Subsequently, definitions of membrane domains and their sizes remain inconsistent. These domains include clathrin-coated pits and lipid rafts, and are believed to compartmentalize the membrane, facilitate the assembly of signaling complexes, and serve as the platforms for response amplification to the extracellular signaling molecules (Kusumi et al., 2012). For instance, it is now estimated that lipid rafts, cholesterol-enriched membrane domains, are predominantly transient complexes of 10-200 nm in size. Such lipid rafts are able to adjust their size and stability in response to ongoing membrane trafficking or signal transduction processes (Eggeling et al., 2009; Jacobson et al., 2019).

An important signaling protein that is present in the plasma membrane of many cell types in vertebrate organisms is H-Ras. The H-Ras protein is a member of the Ras protein family, which consists of small GTPases that activate intracellular signaling cascades, and thereby regulate crucial biological processes taking place in various cells, such as growth, proliferation, and differentiation (Malumbres and Barbacid, 2003). Gain-of-function mutations in genes encoding Ras proteins are found in ca. 25% of human cancers, which makes Ras proteins interesting targets of cancer therapies (Hobbs et al., 2016). These proteins are mainly localized at the plasma membrane, though some fractions have also been reported to be present in membranes of endosomes, the endoplasmic reticulum, and Golgi apparatus. Various Ras isoforms exist, which differ predominantly in their so-called hyper variable region (HVR), formed by 25 amino acids present in their carboxyl-terminal end. The most carboxyl-terminal part of the HVR comprises an anchoring domain, which is responsible for anchoring Ras proteins in the cytoplasmic leaflet of the cell membranes upon posttranslational modifications, mostly the addition of lipid groups. In H-Ras, this domain comprises a CAAX motif, which can be farnesylated, and two cysteine residues that can be (reversibly) palmitoylated (Brunsveld et al., 2009; Willumsen et al., 1984).

Single-molecule microscopy (SMM) has been used in many studies to visualize individual molecules in the plasma membrane to study and characterize their organization and mobility. This technique is mostly performed using advanced fluorescence microscope techniques, such as light sheet fluorescence microscopy (LSFM) or total internal reflection fluorescence microscopy (TIRFM). The used microscopy setups are equipped with a laser for excitation of the fluorescent molecules and with a highly sensitive charged-couple device (CCD) or a complementary metal oxide semiconductor (CMOS) camera for capturing the emitted photons. Molecules subjected to SMM experimentation are often fluorescently labeled. There are several options for selecting a suitable fluorescent label, based on the biological model used or a spectrum of excitation laser light available in the setup (Chudakov et al., 2010; Harms et al., 2001; Seefeldt et al., 2008). Most often, autofluorescent proteins, such as Green or Yellow Fluorescent Proteins (GFP, YFP) fused with an endogenous protein, are used, as they are not toxic to living organisms and their use does not require permeabilization or fixation of the cells.

The mobility of Ras proteins or Ras membrane anchors fused to autofluorescent proteins has been extensively studied and characterized by SMM in cultured cells (Lommerse et al., 2004; Lommerse et al., 2005), with a positional accuracy of up to 30 nm and a temporal resolution in the span from 5 to 50 milliseconds (Harms et al., 1999; Lommerse et al., 2004). Studies in which the H-Ras anchoring domain and full-length H-Ras protein were fused with YFP (referred to as YFP-C10H-Ras and YFP-H-Ras, respectively), showed that the anchor exhibits similar mobility patterns to other anchors of human Ras proteins (e.g. K-Ras), and to the full-length H-Ras protein (Lommerse et al., 2006). It was demonstrated that these protein populations segregate into a slow- and a fast-diffusing fraction. The slow-diffusing fraction of the YFP-fused anchors was found to confine to ca. 200 nm domains, whereas the full-length was only found confined in such domains upon insulin-induced activation and mutation of the protein that induces constitutive activation (YFP-H-Ras^V12^) (Lommerse et al., 2004; Lommerse et al., 2005).

Studying cultured cells has the advantage of easier control and manipulation when compared to living organisms. Nevertheless, studies using cultured cell models do not take into consideration the influence of cell-cell interactions and extracellular stimuli that are present within a tissue. Furthermore, they do not take into account factors an entire organism might be presented with that alter the context of the cells under investigation, such as changes due to the diurnal cycle or the response to a stressor. Therefore, in order to perform SMM studies with more translational value, we extended the applicability of the single-molecule research to a more physiologically relevant system, using the zebrafish embryo as a model organism for studying protein mobility patterns *in vivo*. Zebrafish embryos serve as excellent model organisms for visual analyses of physiological processes, due to their optical clarity (Canedo and Rocha, 2021; Detrich et al., 2011; Garcia et al., 2016; Gore et al., 2018; Lieschke and Currie, 2007). The high fecundity and short generation time of the zebrafish facilitate genetic screens and identification of mutant phenotypes (Haffter et al., 1996; Reisser et al., 2018). Apart from that, genetically modified zebrafish embryos can be readily created by use of microinjection techniques, which makes this organism an excellent model for research using fluorescently labeled cells and proteins.

In a previous study, we used a TIRFM-based approach to perform SMM in zebrafish, and we analyzed the dynamics of YFP-C10H-Ras in epidermal cells of two-day-old embryos. The observed mobility patterns in the zebrafish embryos were different from those found in cultured cells, which underlined the importance of performing this type of studies *in vivo*. Therefore, in the present study, we have extended this application of the *in vivo* SMM technique to the full-length H-Ras protein. In addition to the wild type protein, we have used a constitutively active and inactive H-Ras mutant (H-Ras^V12^ and H-Ras^N17^, respectively) to examine how the protein activity influences the patterns of diffusion and confinement of the H-Ras molecules. Furthermore, we have studied the alterations in the mobility pattern during embryonic development and performed experiments with YFP-C10H-Ras and the full-length H-Ras in human embryonic kidney cells (HEK293T) to compare the results obtained in the zebrafish embryos with results obtained in cultured cells using the same experimental protocol.

Our findings reveal that for YFP-C10H-Ras, YFP-H-Ras, YFP-H-Ras^V12^ and YFP-H-Ras^N17^, in epidermal cells of the zebrafish embryos and in cultured HEK293T cells, a slow- and a fast-diffusing population of molecules can be distinguished and that both populations show confined diffusion. Minor differences in the sizes of the populations, diffusion coefficients and confinement sizes were detected between the proteins and models studied. Interestingly, in zebrafish embryos, the mobility pattern does not change during embryonic development and the variability between individual embryos is smaller than the variability between different areas in the epidermis of one embryo.

## RESULTS

### The mobility pattern of YFP-C10H-Ras and YFP-H-Ras in HEK293T cells

As an initial step in our SMM study, the YFP fusion proteins of the H-Ras anchoring domain and full-length H-Ras protein (YFP-C10H-Ras and YFP-H-Ras respectively) were studied in cultured HEK293T cells using our TIRFM setup. These experiments aimed to compare findings obtained using the TIRFM setup with data previously obtained in cell cultures (Bobroff, 1986; Lommerse et al., 2005), and with our previous findings in zebrafish embryos (Schaaf et al., 2009). Prior to the SMM imaging, cells were transiently transfected with YFP-C10H-Ras and YFP-H-Ras expression vectors and screened in order to analyze the expression levels and sub-cellular localization of the fluorescent proteins. Images of HEK293T cells expressing YFP-C10H-Ras, obtained through confocal laser scanning microscopy, indicated predominant membrane localization of the signal coming from the YFP fused with the C10H-Ras membrane anchor, which is in line with patterns observed before in mouse fibroblast and human embryonic kidney cells (3T3-A14 and ts201, respectively) (Lommerse et al., 2004; Lommerse et al., 2005).

Three days after transfection, YFP-C10H-Ras and YFP-H-Ras expression levels in HEK293T cells were significantly decreased. The sparse distribution allowed for identification of single YFP-C10H-Ras molecules in the basal cell membrane, using our TIRFM setup. Diffraction-limited fluorescence intensity peaks were observed and Gaussian curves were fitted over these peaks (Fig. 1A, B). FWHM and intensity values of the Gaussian distributions corresponding to signals from single YFP molecules were established in fixed HEK293T cells, taking into consideration only these molecules that displayed single-step photobleaching, indicating that their signal originated from an individual YFP molecule. These values were subsequently used to determine cutoff values for peak selection. The signal-to-noise ratio (SNR), defined as the quotient between the average intensity of a single fluorophore (2215 counts) and the standard deviation of the background signal (74 counts), approximated to 30. The positional accuracy equaled approximately 22 nm in one dimension for the localization of these single molecules.

**FIGURE 1.**
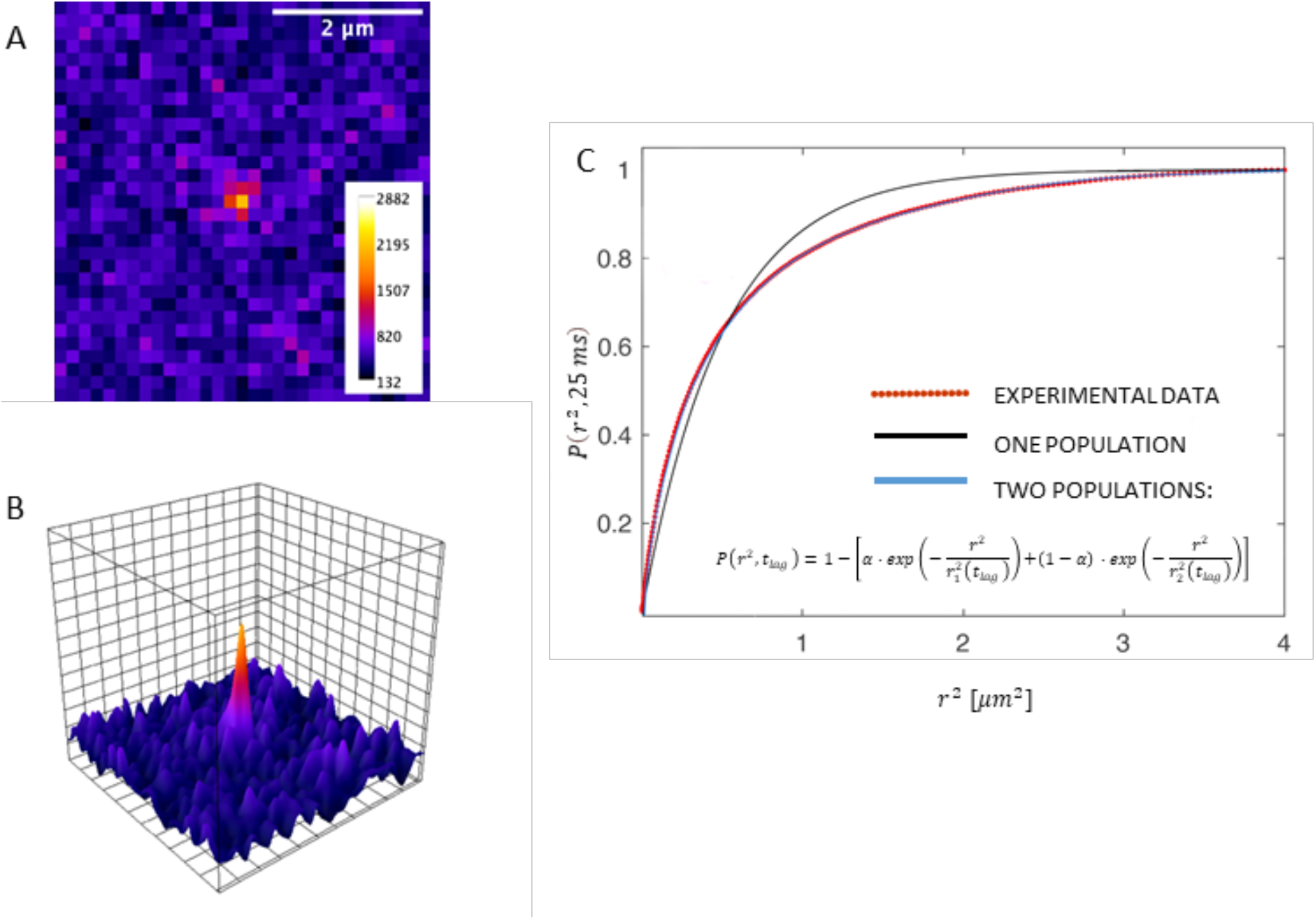
Single-Molecule Microscopy (SMM) analysis of protein localization and mobility. **(A)** SMM intensity map showing signal of a YFP-C10H-Ras molecule located in the basal membrane of a HEK29T cell. **(B)** 3D representation of the image shown in (A), depicting fluorescence intensities of each pixel. One visible intensity peak is shown that can be attributed to a single YFP-C10H-Ras molecule. Over these peaks, two-dimensional Gaussian surfaces were fitted. **(C)** Representative cumulative distribution plot of squared displacements determined using PICS analysis. Data are shown in red, and plots are best using a two populations biexponential model (in blue, formula shown). Fitting of the data points to the two populations model allows for calculation of a relative size of the subpopulations (α) and their mean squared displacements (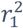 and 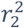). This procedure was repeated for each of the time lags used.

Image sequences were acquired using a time lag of 25 milliseconds, and the mobility pattern of the proteins was determined by Particle Image Correlation Spectroscopy (PICS) software (Semrau and Schmidt, 2007). Using a multistep analysis, information was yielded for five different time lags: 25, 50, 75, 100, and 125 milliseconds. By means of the PICS software, correlations between the location of molecules in consecutive frames were determined, and cumulative probability distributions of the displacements were generated for each time lag. The data were then fitted to a one- or two-population model (Fig. 1C). The two-population model fitted significantly better, indicating the presence of two fractions of molecules with different mobility patterns. Using these curves, the relative size of the fast-diffusing fraction (*α*) and the mean squared displacements of the fast and slow diffusing fractions (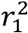 and 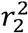) were established for each time lag.

Subsequently, for both YFP-C10H-Ras and YFP-H-Ras, the parameters *α*, 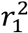, and 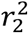 were plotted against the chosen time lags (Fig. 2). The relative size of the fast-diffusing fraction *α* was constant over all time lags, and equaled 77.8 ± 0.4 % and 78.0 ± 0.5 % for YFP-C10H-Ras and YFP-H-Ras, respectively. The curves presenting the relationship between mean squared displacements (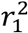, and 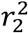) and the time lag for the fast and slow diffusing fractions are non-linear and reach a plateau. We therefore fitted these curves using a confined diffusion model, in which the movement of YFP-C10H-Ras and YFP-H-Ras is confined within an area of size *L* with an initial diffusion coefficient *D*_0_ (Bobroff, 1986; Lingwood and Simons, 2010; Schaaf et al., 2009). The mobility patterns of the two fusion proteins were remarkably similar. For YFP-C10H-Ras, *D*_0_ of the fast-diffusing fraction equaled 1.40 ± 0.20 μm^2^ s^−1^ and the size of the confinement area *L* equaled 480 ± 7 nm. For YFP-H-Ras, the fast-diffusing fraction had a *D*_0_ of 1.22 ± 0.13 μm^2^ s^−1^ and was confined in an area of which *L* equaled 576 ± 11 nm. The slow-diffusing fractions of YFP-C10H-Ras and YFP-H-Ras moved dramatically slower (*D*_0_ of 0.158 ± 0.056 μm^2^ s^−1^ and 0.173 ± 0.036 μm^2^ s^−1^) and were confined in smaller areas (*L* of 157 ± 5 nm and 219 ± 8 nm).

**FIGURE 2.**
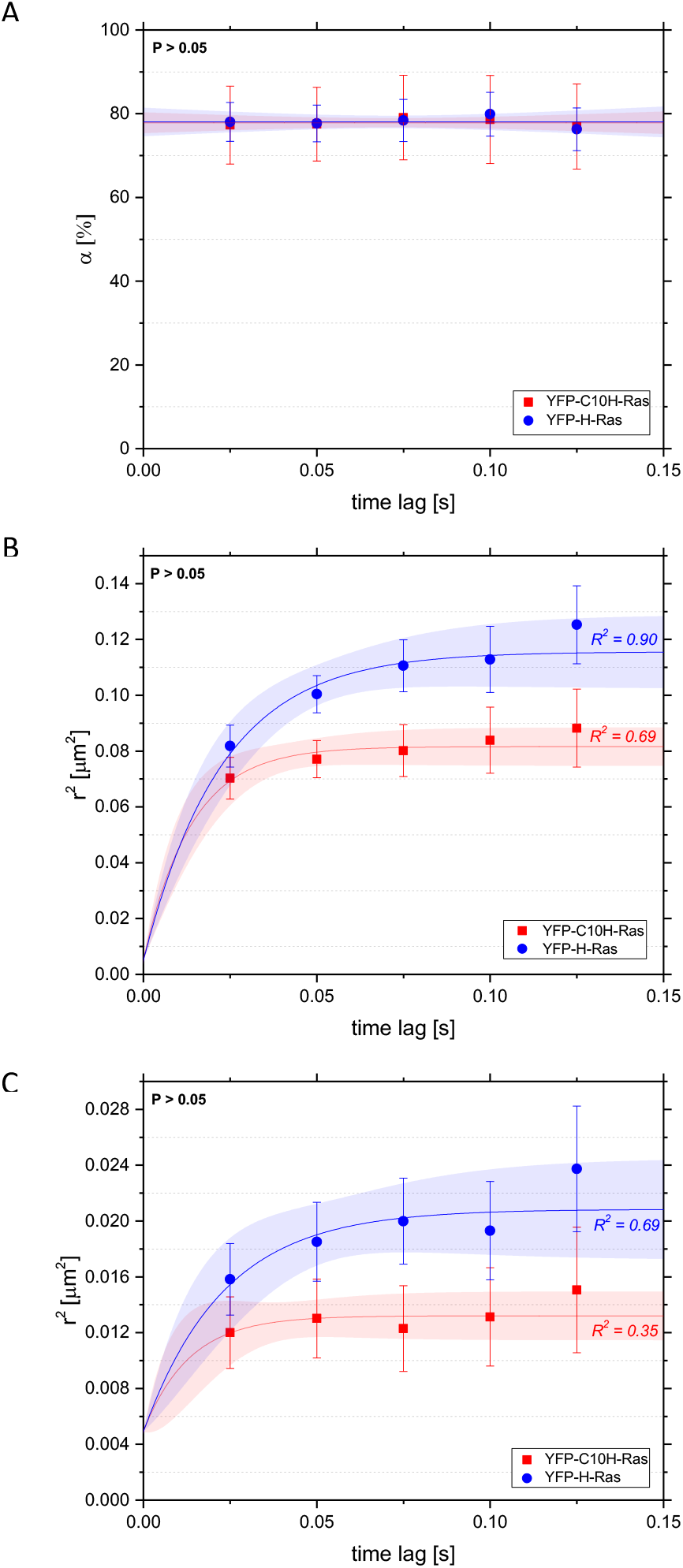
Mobility patterns of YFP-C10H-Ras and YFP-H-Ras in HEK293T cells. (**A**) Fraction size of the fast-diffusing population, plotted against the time lag. (B) Mean squared displacements plotted against the time lag for the fast-diffusing fraction. (C) Mean squared displacements plotted against the time lag for the slow-diffusing fraction. Results of the fits are summarized in Table 1. Statistical analysis was performed using an unpaired t-test. Pearson correlation coefficients R^2^ are presented to show fitness of the data to the model of confined diffusion.

### The mobility pattern of YFP-C10H-Ras and YFP-H-Ras in epidermal cells of zebrafish embryos

In order to successfully image membrane proteins in living zebrafish embryos, we focused our observation on the outer epidermal cell layer in the tail fin of 2 dpf zebrafish embryos (Fig. 3). Embryos were injected with DNA constructs of both YFP-C10H-Ras and YFP-H-Ras at the one-cell stage in order to stably express the YFP-tagged H-Ras anchor and the full-length H-Ras protein in the zebrafish tail fin. This outer cell layer of the skin (the superficial stratum) is a homogenous layer of living cells, and forms the upper part of the skin (the epidermis) together with an underlying cell layer. The homogeneity of this layer was illustrated by confocal microscopic images of 2 dpf embryos of a transgenic zebrafish line expressing GFP-C10H-Ras in all cells.

**FIGURE 3.**
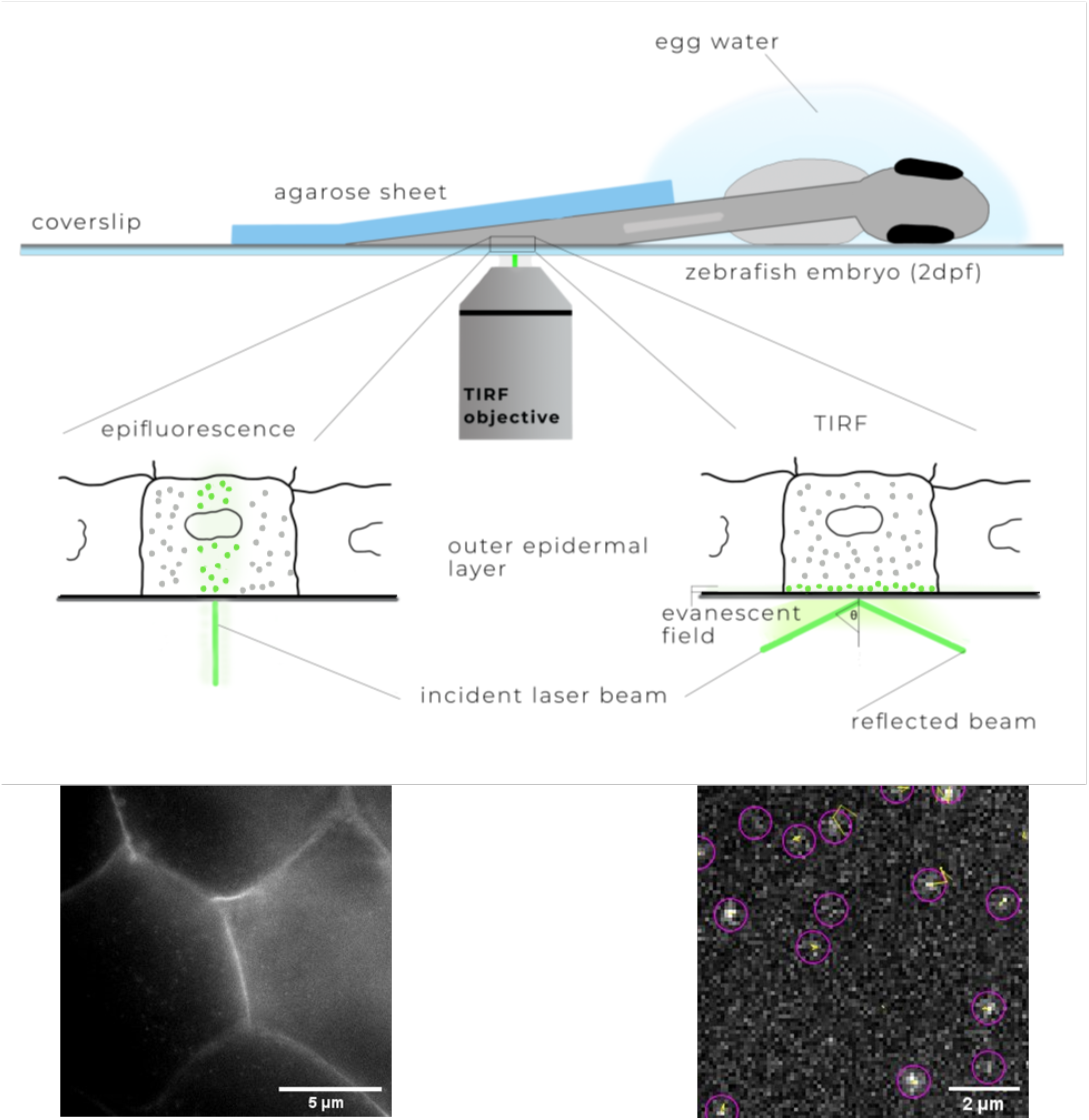
Schematic overview of SMM applied to a living zebrafish embryo. A zebrafish embryo was injected with an expression vector for a YFP fusion protein shortly after fertilization. At 2 dpf, it was placed on a coverslip coated with poly-L-lysine in a drop of egg water. The tail region of the embryo was covered with a 0.75-mm-thick agarose sheet. On the lower left part of the figure, an epifluorescence picture is presented of the outer layer of the epidermis showing the fluorescent signal of YFP-C10-H-Ras in the cell membranes. On the lower right, a TIRFM image from the same FOV is presented, with examples of individual YFP molecules shown in circles and their trajectories in yellow.

The tail fins of 2 dpf zebrafish embryos are morphologically stable enough to resist coverage with a 0.75-mm-thick sheet of agarose, which was used in order to gently press the tail fin towards the surface. This enables the evanescent field to excite the fluorophores present in the outer membrane of cells in the outer epidermal cell layer. For imaging, we focused on the tail fin region of the zebrafish embryo, while the rest of the zebrafish body was immersed in a drop of water. Zebrafish vital functions, such as heartbeat and the blood flow in the cardiovascular system, were checked under a stereofluorescence microscope post-imaging.

As in HEK293T cells, we observed fast- and slow-diffusing protein fractions in zebrafish embryos (Fig. 4). The fast-diffusing fraction size *α* equaled 65.0 ± 0.5 % for the YFP-C10H-Ras, and 65.9 ± 0.6 % for YFP-H-Ras. In both of the fractions, molecules followed a confined diffusion pattern, with sizes of the confinement areas *L* for the fast-diffusing fractions being approximately 3-4 times larger than for the slow-diffusing ones (See Table 1, part 1.1, 1.2). The initial diffusion coefficient *D*_0_ of the fast-diffusing fraction equaled 1.10 ± 0.15 μm^2^ s^−1^ for YFP-C10H-Ras and 0.71 ± 0.05 μm^2^ s^−1^ for YFP-H-Ras. The initial diffusion coefficients for the slow-diffusing fractions were at least 8 times lower than the ones for their fast-diffusing counterparts. Overall, the results show that there is not a significant difference between HEK293T and zebrafish embryo models, where we determine the mobility patters of YFP-H-Ras and YFP-C10H-Ras, with a notable exception of the initial diffusion coefficients *D*_0_ for YFP-H-Ras, which, both in the case of the fast- and the slow-diffusing fraction, were significantly lower in the zebrafish embryo model (See Table 1).

**FIGURE 4.**
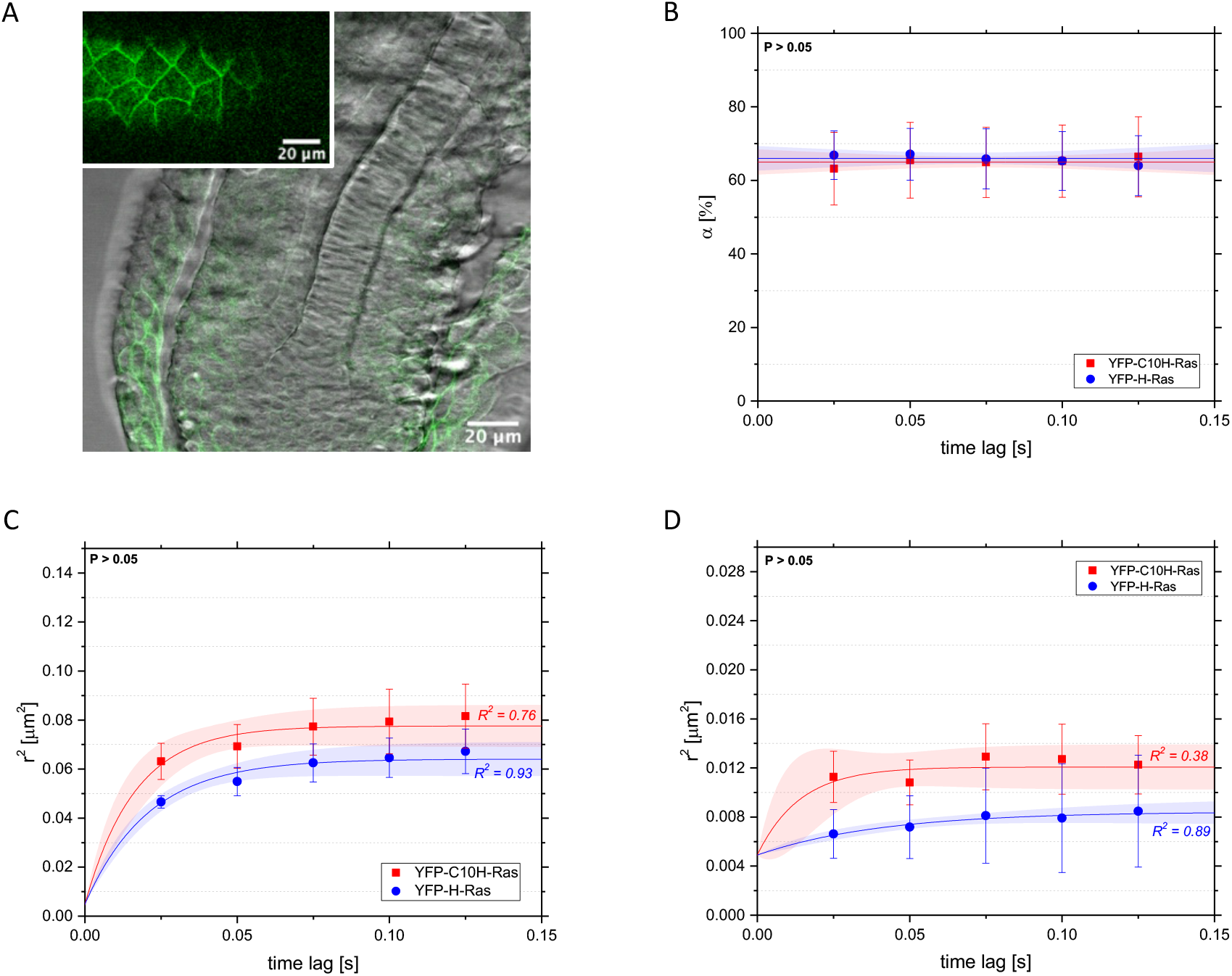
Mobility patterns of YFP-C10H-Ras and YFP-H-Ras in epidermal cells of the zebrafish embryos. **(A)** Overlay of brightfield and fluorescence microscopy image of the zebrafish embryo epidermal cell layer expressing GFP-C10H-Ras at 2 dpf (GFP-C10H-Ras transgenic line). The image shows that the embryonic epidermis consists of a homogenous cell population. Inset shows fluorescence microscopy image depicting a part of the zebrafish tail fin membrane. (**B**) Fraction size of the fast-diffusing population, plotted against the time lag. **(C)** Mean squared displacements plotted against the time lag for the fast-diffusing fraction. **(D)** Mean squared displacements plotted against the time lag for the slow-diffusing fraction. Results of the fits are summarized in Table 1. Statistical analysis was performed using an unpaired t-test. Pearson correlation coefficients R^2^ are presented to show fitness of the data to the model of confined diffusion.

**TABLE 1.** Results of the analysis of the mobility patterns of YFP-C10H-Ras, YFP-H-Ras, YFP-H-Ras^V12^, YFP-H-Ras^N1^

| Experiment | $D_{\text{OFD}} [\mu\text{m}^2 \text{s}^{-1}]$ | $D_{\text{OSD}} [\mu\text{m}^2 \text{s}^{-1}]$ | $L_{\text{FD}} [\text{nm}]$ | $L_{\text{SD}} [\text{nm}]$ | $\alpha$ [%] |
| --- | --- | --- | --- | --- | --- |
| **YFP-C10H-Ras: comparison between HEK293T cells and zebrafish embryos** |  |  |  |  |  |
| HEK293T | $1.40 \pm 0.20$ | $0.158 \pm 0.056$ | $480 \pm 7$ | $157 \pm 5$ | $77.8 \pm 0.4$ |
| zebrafish | $1.10 \pm 0.15$ | $0.128 \pm 0.057$ | $467 \pm 9$ | $147 \pm 6$ | $65.0 \pm 0.5$ |
| **YFP-H-Ras: comparison between HEK293T cells and zebrafish embryos** |  |  |  |  |  |
| HEK293T | $1.22 \pm 0.13$ | $0.173 \pm 0.036$ | $576 \pm 11$ | $219 \pm 8$ | $78.0 \pm 0.5$ |
| zebrafish* | $0.71 \pm 0.05$ | $0.022 \pm 0.003$ | $421 \pm 8$ | $103 \pm 5$ | $65.9 \pm 0.6$ |
| **YFP-C10H-Ras in zebrafish embryos: developmental study** |  |  |  |  |  |
| 48 hpf | $1.84 \pm 0.44$ | $0.114 \pm 0.036$ | $507 \pm 10$ | $170 \pm 7$ | $68.2 \pm 1.3$ |
| 56 hpf | $1.54 \pm 0.07$ | $0.199 \pm 0.047$ | $610 \pm 5$ | $197 \pm 6$ | $60.6 \pm 0.5$ |
| 72 hpf | $1.27 \pm 0.04$ | $0.128 \pm 0.032$ | $616 \pm 3$ | $124 \pm 2$ | $62.9 \pm 0.4$ |
| 80 hpf | $1.10 \pm 0.20$ | $0.107 \pm 0.026$ | $572 \pm 19$ | $153 \pm 3$ | $52.4 \pm 0.4$ |
| **YFP-H-Ras<sup>(WT, N17, V12)</sup> in zebrafish embryos: comparison of the mutants** |  |  |  |  |  |
| YFP-H-Ras | $0.71 \pm 0.05$ | $0.022 \pm 0.003$ | $421 \pm 8$ | $103 \pm 5$ | $65.9 \pm 0.6$ |
| YFP-H-Ras<sup>N17</sup> | $0.72 \pm 0.07$ | $0.030 \pm 0.002$ | $444 \pm 9$ | $100 \pm 2$ | $63.2 \pm 0.1$ |
| YFP-H-Ras<sup>V12</sup> | $1.05 \pm 0.06$ | $0.055 \pm 0.006$ | $572 \pm 7$ | $132 \pm 5$ | $61.8 \pm 0.8$ |

\*Results of YFP-H-Ras used for comparison between the models are taken from experiment in which YFP-H-Ras, YFP-H-Ras<sup>V12</sup> and YFP-H-Ras<sup>N17</sup> were studied.

### The mobility pattern of YFP-C10H-Ras in zebrafish embryos at different developmental stages

To study the stability of the data obtained in the zebrafish embryos over different developmental stages, we analyzed the mobility patterns of YFP-C10H-Ras at 48, 56, 72 and 80 hpf. At each stage, the values for the parameters *α*, 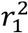, and 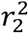 were determined for each time lag, and *D*_0_ and *L* were determined for both fast- and slow-diffusing populations (Fig. 5). The results showed that a difference in the developmental stage of the zebrafish embryo did not significantly influence any of these parameters (Table 2). This means that the C10H-Ras mobility patterns remain stable within the time frame of our experiments.

**FIGURE 5.**
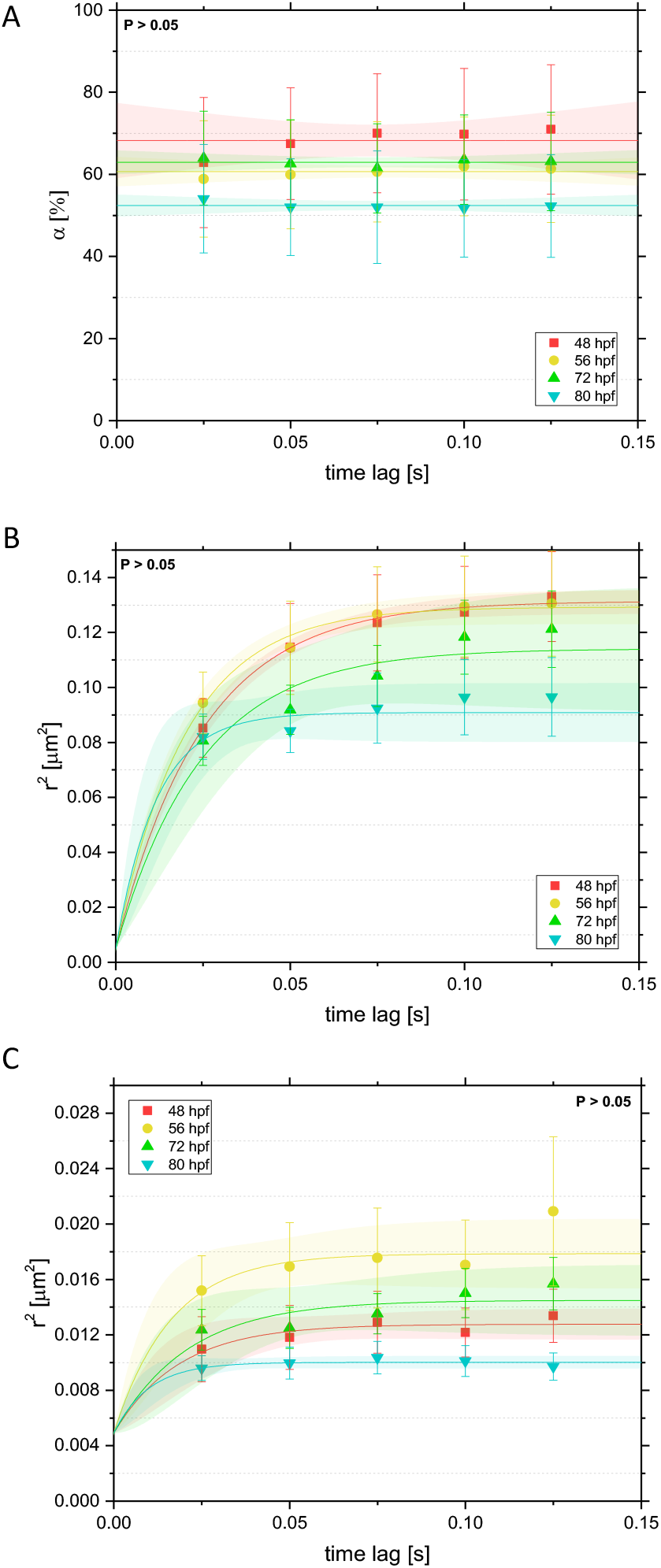
Mobility patterns of YFP-C10H-Ras in epidermal cells of the zebrafish embryos at different developmental stages. (**A**) Fraction size of the fast-diffusing population, plotted against the time lag. **(B)** Mean squared displacements plotted against the time lag for the fast-diffusing fraction. **(C)** Mean squared displacements plotted against the time lag for the slow-diffusing fraction. Results of the fits are summarized in Table 1. Statistical analysis was performed, using a one-way ANOVA.

**TABLE 2.**
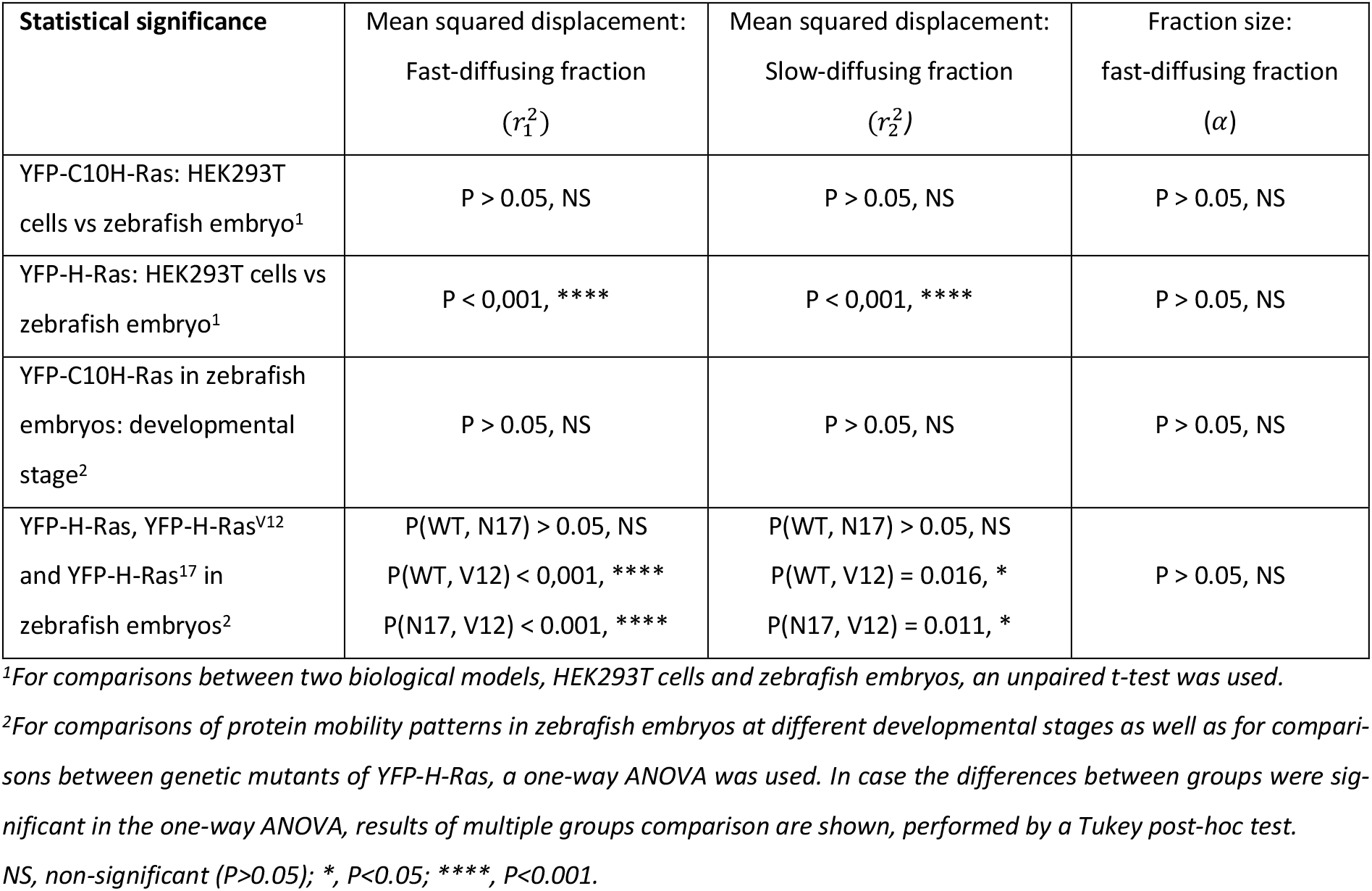
Statistical analysis performed for the values of mean squared displacements and fast-diffusing fraction sizes obtained experimentally for the 25 ms time lag.

### The mobility pattern of YFP-H-Ras^V12^ and YFP-H-Ras^N17^ in epidermal cells of zebrafish embryos

To study the effect of the activation state of the H-Ras protein, the constitutively active H-Ras^V12^ mutant (with a valine replacing a glycine at position 12) and the inactive mutant H-Ras^N17^ (with an asparagine replacing a serine at position 17) were used. The fusion proteins YFP-H-Ras, YFP-H-Ras^V12^ and YFP-H-Ras^N17^ were expressed in 2 dpf zebrafish embryos, and the mobility patterns of the each of these constructs were analyzed. Again, we observed a fast- and a slow-diffusing fractions of molecules. The size of the fast-diffusing fraction did not differ significantly between the constructs, and equaled 66.0 ± 0.5 % for YFP-H-Ras, 61.7 ± 0.7 % for YFP-H-Ras^V12^, and 63.2 ± 0.2 % for YFP-H-Ras^N17^ (Fig. 6A). All fractions showed confined diffusion. Initial diffusion coefficients *D*_0_ for the fast-diffusing fraction equaled 0.70 ± 0.05 μm^2^ s^−1^ for YFP-H-Ras, 1.05 ± 0.06 μm^2^ s^−1^ for YFP-H-Ras^V12^, and 0.72 ± 0.07 μm^2^ s^−1^ for YFP-H-Ras^N17^, with the diffusion coefficient for YFP-H-Ras^V12^ being significantly higher than that for the other two constructs (Fig. 6B). Similarly, the size of the confinement area *L* for YFP-H-Ras^V12^ (572 ± 7 nm) was significantly higher than those for YFP-H-Ras and YFP-H-Ras^N17^ (421 ± 8 nm and 444 ± 9 nm, respectively). The initial diffusion coefficients and the sizes of the confinement areas for the slow diffusing fractions of YFP-H-Ras^V12^ and YFP-H-Ras^N17^ did not show any significant difference from the values determined for the wild type YFP-H-Ras (0.022 ± 0.003 μm^2^ s^−1^ and 103 ± 5 nm; for detailed results, see Table 1, Table 2, and Fig. 6C).

**FIGURE 6:**
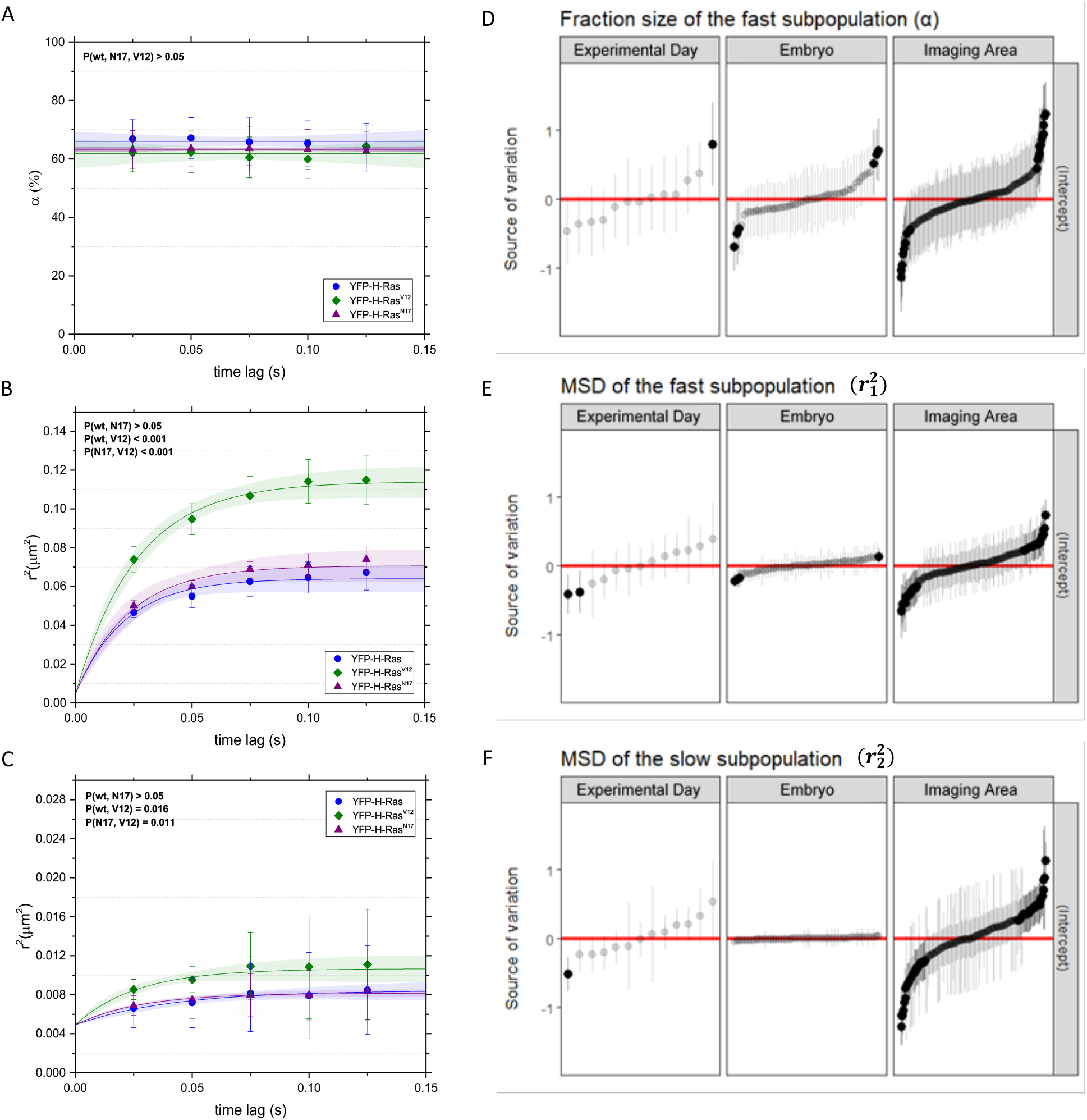
Mobility patterns of YFP-H-Ras and its mutants, YFP-H-Ras^V12^ and YFP-H-Ras^N17^, in epidermal cells of the zebrafish embryos at different developmental stages. (**A**) Fraction size of the fast-diffusing population, plotted against the time lag. **(B)** Mean squared displacements plotted against the time lag for the fast-diffusing fraction. (C) Mean squared displacements plotted against the time lag for the slow-diffusing fraction. Results of the fits are summarized in Table 1. Statistical analysis was performed, using a one-way ANOVA. **(D, E, F)** Caterpillar plots presenting the effect range of a different experimental day, different embryos within an experimental day, and different areas within an embryo to an overall variability in the fast-diffusing fraction size **(D)**, mean squared displacement of the fast-diffusing fraction **(E)**, and slow-diffusing fraction **(F)**. Effect ranges represent the relative deviation of group intercepts from the overall mean with 95% confidence intervals. Red lines indicate the overall mean of the data, while black points signify groups that significantly deviate from the overall mean among batches, individual embryos, and areas. The data points are sorted based on their deviation from the total average, with the ones most negatively deviating placed on the left. The values in **(D)** were logit-transformed and the values in **(E)** and **(F)** were logarithm-transformed, to meet the statistical hypotheses of the hierarchical linear model.

### The sources of variability of the results

Finally, we analyzed the sources of variability of the results obtained in zebrafish embryos for YFP-H-Ras and the two mutants, YFP-H-Ras^V12^ and YFP-H-Ras^N17^. For this purpose, we first studied the correlation between the number of individual molecules within images and the parameters *α*, 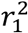, and 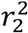. Correlation coefficients between the number of molecules and the corresponding parameters were low (0.22, 0.19 and 0.28, respectively), indicating that the correlation between the number of molecules in an image and the parameters describing protein mobility is very weak. Thus, we affirm that the number of molecules has a negligible impact on the H-Ras mobility analysis.

Secondly, the contribution of three factors to the overall variability in the results was analyzed. These factors were: the different batches of zebrafish embryos, different individual embryos, and the different imaged areas. By use of hierarchical linear mixed models, we generated caterpillar plots for *α*, 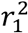, and 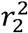, which present random effects distribution, being the deviation of the group intercept from an overall mean, and the contribution of each of the three factors to the overall mean of a given parameter (Fig. 6D-F). Subsequently, we quantified the percentage contribution of each of the selected sources of variation towards the overall data variability (Table 3). These caterpillar plots and the quantitative source of variation analysis both show that, for every parameter, most of the random effects come from imaging embryos on different experimental days (contribution to the total variability of 33.1% for the fraction size, 49.2% for the fast-diffusing fraction diffusion coefficient) and imaging different areas within the epidermis of the same zebrafish embryo (contribution of 50.1% for the slow-diffusing subpopulation diffusion coefficient). Interestingly, imaging different embryos from the same batch was the smallest source of variability for all parameters, and did not introduce nearly as much variability as imaging of different areas within the same individual zebrafish embryo (Table 3).

**TABLE 3.** Summary of the variance values (σ^2^) per source parameter and their percentage contribution (%) towards the total variance for the given parameter.

| Source of variation | $\alpha$ | | $r_1^2$ | | $r_2^2$ | |
| --- | --- | --- | --- | --- | --- | --- |
| | $\sigma^2$ | % | $\sigma^2$ | % | $\sigma^2$ | % |
| Experimental Day | 0.188 | 33.1 | 0.097 | 49.2 | 0.111 | 35.6 |
| Embryo | 0.122 | 21.5 | 0.018 | 9.0 | 0.004 | 1.4 |
| Imaging Area | 0.173 | 30.4 | 0.066 | 33.7 | 0.157 | 50.5 |
| Residual<sup>1</sup> | 0.085 | 14.9 | 0.016 | 8.1 | 0.039 | 12.5 |

<sup>1</sup>Residuals account for variance that cannot be explained by the experimental design.

## DISCUSSION

In the present study, we have applied single-molecule microscopy to an *in vivo* model system, using living zebrafish embryos. As our model molecule we have used a YFP-fusion of H-Ras, a signaling protein that is anchored in the cytoplasmic leaflet of the plasma membrane. Based on the observed mobility patterns in epidermal cells of the embryos, two fractions of H-Ras molecules were distinguished: a fast- and a slow-diffusing one, which both show confined diffusion. The fast fraction contains the majority of molecules and shows an initial diffusion coefficient that is approximately 10 to 15 times higher than that of the slow-diffusing fraction. Similar mobility patterns were obtained when only the membrane anchor of the H-Ras protein was fused to YFP, or when a constitutively inactive H-Ras mutant (H-Ras^N17^) was studied. A constitutively active H-Ras mutant (H-Ras^V17^), however, showed an increased mobility with a higher diffusion coefficient and a larger confinement area. Interestingly, the largest variation in these experiments was not found between individual embryos, but between different experimental days and the areas within a single embryo.

In this study, the observation of a fast and a slow diffusing fraction of H-Ras molecules that were both confined was highly consistent between experimental models used, i.e., between cultured human embryonic kidney (HEK293T) cells and epidermal cells of zebrafish embryos, between different stages of zebrafish embryonic development, and between the full length H-Ras protein, the constitutively active and inactive mutant and the H-Ras membrane anchor (C10H-Ras). These fast and slow diffusing fractions have been found before in several single-molecule microscopy studies on H-Ras or C10H-Ras (the latter also in zebrafish embryos), with diffusion coefficients comparable to those found in the present study (Lommerse et al., 2004; Lommerse et al., 2005; Lommerse et al., 2006; Schaaf et al., 2009). However, confinement of these fractions was not always observed, which in most cases may have been due to the shorter time ranges used in these studies combined with larger variation or smaller sample sizes.

Although the observed mobility patterns were remarkably stable in our experiments, we did observe two cases of altered mobility. First, the full-length H-Ras protein displayed a lower diffusion coefficient in the epidermal cells of the embryos compared to the diffusion coefficients found in the HEK293T cells. This difference was observed for both fractions of this protein, but was not found for any of the fractions of the C10H-Ras protein. Second, the constitutively active H-Ras mutant protein H-Ras^V17^ showed increased mobility of its fast-diffusing fraction compared to the wild type H-Ras protein, which was reflected in an increased initial diffusion coefficient and a larger confinement area compared to the wild type H-Ras protein. In a previous study on the mobility of this mutant in cultured mouse embryonic fibroblasts (3T3 cells), this increased mobility was not observed (Lommerse et al., 2006). In addition to the shorter time range, larger variation and smaller sample sizes mentioned earlier as an explanation for such discrepancies, the cell type used may be also be a possible factor underlying the absence of this effect in the earlier study.

Using the mobility data obtained in zebrafish embryos, we studied the factors that contributed to the variation in these data. Interestingly, differences between individual embryos only had a modest effect on the overall variation in the results. However, different experimental days and different areas of the embryonic epidermis that we imaged both appeared to be factors that showed large contribution to the overall variation in the data. The area that we imaged had a size of 16.6×16.6 μm, which is approximately the area of a single cell (see Fig. 3), so we assume that cell-to-cell variation within the epidermal tissue was a main source of the observed variation. This indicates that the cellular context determines H-Ras mobility, and that differences between cells within the epidermal tissue of one individual embryo are larger than the differences in average cellular context in the tissue between embryos. The observation that the membrane anchor of H-Ras fused to YFP (C10H-Ras) shows a mobility pattern similar to the full length YFP-H-Ras fusion protein indicates that the functionality of the protein does not play a role and that therefore interactions with structural elements of the cell, such as specific membrane domains and the cytoskeleton, are determining the dynamics of these proteins. In addition to the cell-to-cell variation, day-to-day variation appeared to contribute largely to the total variation. We assert that these differences may for a large part originate from an inherent variability between different zebrafish batches. The AB/TL zebrafish wild type strain that was used is not an inbred strain, and its genetic heterogeneity might underlie batch to batch variation. Furthermore, small differences in treatment conditions might affect embryonic development, alter complex metabolic pathways, and induce stress responses, and thereby cause, to some extent, the observed differences in the values of the measured dynamic parameters on different experimental days.

Lipid-tethered Ras protein family members have been reported to possess an intrinsic ability to form nanoclusters composed by a small number of Ras molecules. It has been specifically shown that H-Ras is localized in largely separated nanoclusters of less than 30 nm that have an average lifetime of less than 1 second (Pezzarossa et al., 2015; Plowman et al., 2005; Prior et al., 2003; Rotblat et al., 2004). It has been suggested that these clusters of H-Ras proteins are localized in lipid rafts, which are thought to be cholesterol-enriched domains in the plasma membrane from 10 to 100 nm in size (Garcia-Parajo et al., 2014; Lingwood and Simons, 2010; Nickels et al., 2015). Due to the selective partitioning of rafts and the surrounding areas, rafts are responsible for controlling specific downstream signaling processes, by enhancing certain protein-protein interactions, while suppressing others. These relatively immobile H-Ras nanoclusters are a result of temporal trapping within these domains and constitute approximately 40% of the total protein population, whereas the rest of the H-Ras molecules remains monomeric and laterally mobile within the plasma membrane (Dadke and Chernoff, 2003; Murakoshi et al., 2004; Rotblat et al., 2004). In the current study, the fraction size of the fast-diffusing H-Ras in living zebrafish embryos equaled approximately 35%, suggesting that this fraction reflects the clustering subpopulation of H-Ras molecules. Thus, H-Ras compartmentalization into confinement areas within the plasma membrane is believed to be necessary to aggregate a high number of H-Ras molecules in order to confine a cytoplasmic signal to an area in which an extracellular signal is received (Simons and Gerl, 2010).

The localization of H-Ras in lipid rafts appears to be enhanced by the presence of the lipid anchor (Pezzarossa et al., 2015; Rotblat et al., 2004) and in our study lipid anchor of H-Ras indeed appears to be sufficient for the formation of the slow-diffusing fraction. In contrast, it has been shown that GTP loading of the N-terminus of the protein decreases the affinity of H-Ras for these microdomains (Rotblat et al., 2004). Relatively high diffusion coefficients have been observed for the H-Ras^V12^ fast-diffusing fractions, which is in line with our results for this mutant. The confinement of the fast-diffusing subpopulation may result from exclusion from lipid rafts, but other cellular structures, such as the actin cytoskeleton, may play an equally important role in confinement of membrane molecules. Multiple studies have suggested that confinement areas can also be formed by impermeable barriers constituted by cytoskeleton reformation. The membrane-associated cytoskeleton is a highly organized structure that is able to create multimolecular membrane-cytoskeleton assemblies, including focal adhesions and podosomes (Garcia-Parajo et al., 2014).

In conclusion, in the present study we have used a previously developed TIRFM-based approach to perform SMM in an intact vertebrate organism. Our study confirms that this technology is highly useful for studying the molecular behavior of individual receptors and signaling molecules, and enables further *in vivo* studies to unravel exact molecular mechanisms governing molecular interactions and their role in physiological and pathological processes, such as skin cancer, wound healing, or tissue regeneration (Kaufman et al., 2016; Kawakami et al., 2004). We have studied the mobility pattern of H-Ras in the epidermis of living zebrafish embryos, and our data show the consistent presence of a fast and a slow diffusing fraction of H-Ras molecules, which both show confinement. Interestingly, the epidermal tissue shows a large degree of heterogeneity between individual cells, and variation in the structural context of cells within this tissue appears to be a main determinant of H-Ras dynamics.

## MATERIAL AND METHODS

### Zebrafish

Wild type ABTL zebrafish (*Danio rerio*) were grown and maintained according to standard protocols (http://ZFIN.org), exposed to a 14h light and 10h dark diurnal cycle at 28°C. Fertilization was performed by natural spawning at the beginning of the light period. Eggs were collected and raised in egg water (60 μg/ml Instant Ocean sea salts, Cincinnati, OH, USA) at 28°C. All experiments performed on living zebrafish embryos were done in compliance with the directives of the local animal welfare committee of Leiden University.

### Cell cultures, transfection, and fixation

In all cell culture experiments, human embryonic kidney cells (HEK293T) were used. Cells were cultured in DMEM (Dulbecco’s Modified Eagle Medium, Invitrogen, Waltham, MA, USA) supplemented with penicillin and streptomycin (10 μg ml^−1^, Invitrogen), Glutamax (10 μg ml^−1^, Invitrogen) and 10% fetal calf serum (Invitrogen) at 37°C in a humidified atmosphere containing 5% of CO2. Cells were passaged every 3 to 4 days and kept in use for a maximum of 12 passages. Before transfection, cells were transferred onto a sterile, glass coverslip (diameter 25 mm, Marienfeld, Lauda-Königshofen, Germany) placed in a well of a 6-well plate. At a confluency level of 20-30%, cells were transfected with 1 μg of DNA per well, using FuGENE 6, according to the manufacturer’s protocol (Roche Molecular Biochemicals, Indianapolis, USA). The transfection efficiency, determined by fluorescence microscopy screening at 48h after transfection (EVOS M7000 Cell Imaging Systems, ThermoScientific, Waltham, MA, USA), was approximately 30%. Cells were imaged at least 4 days post-transfection to lower the expression level of the YFP-fused proteins and to be able to efficiently observe single molecules. Prior to the single-molecule microscopy experiments, DMEM was replaced by PBS (phosphate-buffered saline; 150 mM NaCl, 10 mM Na_2_HPO_4_/NaH_2_PO_4_, pH 7.4) kept at room temperature. Finally, the coverslips with cells were mounted on a microscope holder, and 500 μl of room-temperature PBS was pipetted onto the cells. For single-step photobleaching experiments, transfected cells were fixed using 4% paraformaldehyde (PFA) in PBS overnight, at 4°C. Immediately before the experiments, the cells were washed three times, 5 minutes each time, with PBS. Finally, the cells were rinsed twice with PBST (PBS with Tween^™^-20, 0.1%), for 5 minutes each time.

### Microinjection of DNA in zebrafish embryos

The cDNAs encoding YFP-C10H-Ras, YFP-H-Ras, YFP-H-Ras^V12^, and YFP-H-Ras^N17^ were cloned from pcDNA3.1(+) mammalian expression plasmids into pCS2(+) plasmids, which then served as vectors for expression of the fluorescent proteins of interest. DNA plasmid microinjections were performed at concentration dose of 30 pg per embryo at the 1-2 cell stage, resulting in a mosaic expression of the fluorescent protein in the zebrafish embryos at later stages. Microinjections were done using a Femtojet microinjector (Eppendorf, Hamburg, Germany) and a micromanipulator with pulled microcapillary pipettes. The procedure of microinjections was controlled under a stereomicroscope (M165C, Leica Microsystems, Wetzlar, Germany). Injected eggs were then left to develop in an incubator at 28°C. Viability and development of the eggs after microinjections was checked on a daily basis using fluorescence stereo- or confocal microscopy.

### Fluorescence stereomicroscopy

For screening zebrafish embryos expressing YFP-fused proteins, a Leica M205FA fluorescence stereomicroscope (Leica Microsystems) was used. Images of the zebrafish embryos were taken using a Leica DFC 345FX camera.

### Confocal laser-scanning microscopy

A Leica SPE confocal laser-scanning microscope (Leica Microsystems) was used to investigate the fluorescent signal in the zebrafish embryo. Excitation was done using an argon laser at 514 nm. Images were obtained using a 20x and 40x non-immersion objectives (NA 0.70, NA 0.80, respectively), and a 63x water immersion objective (NA 1.20).

### Total internal reflection fluorescence microscopy (TIRFM)

Glass coverslips were washed with 99% ethanol (twice), HPLC-grade water (twice), KOH (1M, twice), and acetone (99%, thrice). Each wash was followed by a 30-minute-long sonication period at 50°C. If not used immediately, the coverslips were stored in acetone. Prior to the mounting of the zebrafish embryos, the glass coverslips were coated with 50 μg ml^−1^ of poly-L-lysine (Sigma-Aldrich, St. Louis, MO, USA) for 5 minutes, followed by a double wash with deionized water and drying with nitrogen gas. Two-day-old zebrafish embryos were equilibrated at room temperature for an hour, anaesthetized with 0.02% aminobenzoic acid ethyl ester (tricaine, Sigma-Aldrich) and dechorionated using tweezers. Subsequently, a single zebrafish embryo was placed on a coverslip with a lateral side against the coverslip, while excess water was aspirated. The tail of the embryo was pressed against the surface of the coverslip by a thin agarose sheet (2%, thickness 0.75 mm). A drop of egg water was added to cover the rest of the embryo’s body. The coverslip with the embryo was placed on a microscope holder and mounted on the TIRFM setup. This setup was a custom-made microscope with a 100x oil-immersion objective (NA 1.45, Nikon, Tokyo, Japan). Excitation was performed using a 515 nm laser (iChrome MLE, Toptica Photonics, Germany), the field of view was set to a 100×100 pixels region with a pixel size of 166 nm, and the laser power equaled to 20% of the maximal laser power (40 mW). The incident laser beam was set at the critical angle against the coverslip-water interface, thus being totally reflected and creating the evanescent wave for excitation of fluorophores close to the coverslip-sample interface. Emission light was filtered using a long pass filter (ET5701p, Chroma Technology, VT, USA), and image sequences were collected using an on-chip multiplication gain CCD camera (model 512B, Cascade, Roper Scientific, Tucson, AZ, USA). Each image sequence contained 1200 frames separated by a 25 ms time lag, resulting in the total acquisition time of 30 seconds.

### Analysis of protein diffusion patterns

The analysis of the position of individual molecules was done as described previously. Fluorescence intensity signals corresponding to YFP molecules were fitted to a two-dimensional Gaussian surface, using previously custom-developed software (Groeneweg et al., 2014; Lommerse et al., 2004; Lommerse et al., 2005). Subsequently, the corresponding signals were filtered based on their full width at half maximum (FWHM) value and their intensity count, based on the microscopy setup. The FWHM and intensity threshold values were obtained in the single-step photobleaching experiments (performed using fixed HEK293T cells), by averaging the Gaussian distributions of 20 different YFP molecules in the last step prior to the photobleaching-induced final intensity drop (using TrackMate plug-in, ImageJ). The location of a molecule was defined by the center of the Gaussian curve. Positional accuracy (*dx*) of the peak localization equaled approximately 22 nm (Groeneweg et al., 2014; Schütz et al., 1997).

To study the mobility pattern of the proteins, Particle Image Correlation Spectroscopy (PICS) software was used, which has been previously described (Semrau and Schmidt, 2007). A multistep analysis was performed for each image sequence acquired, yielding information for five different time lags of 25, 50, 75, 100, and 125 milliseconds. In PICS analysis, individual particles are not tracked, but correlations between the location of molecules in consecutive frames are determined. This way, cumulative probability distributions of the squared displacements were generated for each of the time lags, and fitted to one- or two-population model. The former is described by the equation:

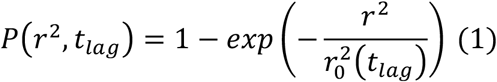

which describes the probability that a particle exhibiting Brownian motion at the arbitrary origin is found within a circle of a radius *r* at the time lag *t_lag_*, and its mean square displacement equals 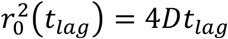. However, in case that the populations of the molecules can be differentiated into two different populations equation (1) is transformed into:

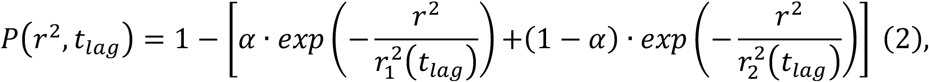

where the mean square displacement of fast-diffusing and slow-diffusing populations are denoted by 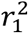 and 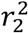, and their relative sizes by *α* and 1 – *α*, respectively (Schaaf et al., 2009). Hence, *α* represents the fraction size of the fast-diffusing H-Ras molecules and is represented as a percentage of the total population.

To examine whether any of these populations confine to a certain area, the values of 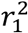 and 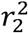 were plotted against the time lag. For the larger time lags the variance increases due to the smaller number of statistically independent measurements of *r*^2^ (Kusumi et al., 1993). The positional accuracy (*dx*) led to a constant offset in *r*^2^of 4 · (*dx*)^2^, which, in our case, equaled 0.0049 μm^2^. The plots were fitted by a free Brownian diffusion model, with a diffusion coefficient *D* in the fitted equation 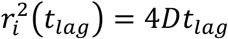, or by a confined diffusion model described by the equation:

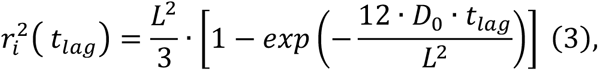

in which the molecules move freely with an initial diffusion coefficient *D*_0_, but are confined by impermeable barriers within an area described by a square of a side length *L*.

### Experimental design

In every experiment, three different zebrafish embryos were selected for the TIRF microscopy imaging. In each of the selected embryos, at least three separate areas were imaged. The data shown reflect a minimum of three independent experiments, each performed on three different days. Six independent experiments were done for YFP-C10H-Ras (18 individual embryos), and three for the wild-type YFP-H-Ras, YFP-H-Ras^N17^, and YFP-H-Ras^V12^ (9 individual embryos per H-Ras construct). For studying the influence of the developmental stage on the YFP-C10H-Ras dynamics, an analogous design was implemented (3 experiments, 3 individual embryos per each experiment), yet the embryos were selected and imaged at 48-, 56-, 72-, and 80-hour post fertilization. In the case of transfected HEK293T cells, three independent experiments were performed for both YFP-C10H-Ras and wild-type YFP-H-Ras. In each of those experiments, three different coverslips with growing cells were selected from a 6-well-plate, and at least three different cells were imaged on each of the coverslips.

### Statistical Analysis

Values of the fast-diffusing population sizes and squared displacements were averaged per experimental day for each time lag in all individual groups (i.e., YFP-C10H-Ras, YFP-H-Ras, YFP-H-Ras^N17^, and YFP-H-Ras^V12^). Statistical analysis of the data (presented in Table 2 for the 25 ms time lag) was performed for each time lag by comparing: (a) results obtained from zebrafish embryos expressing wild-type YFP-H-Ras, YFP-H-Ras^N17^, and YFP-H-Ras^V12^; (b) results obtained for HEK293T cells expressing YFP-C10H-Ras and wild-type YFP-H-Ras, (c) results obtained zebrafish embryos expressing YFP-C10H-Ras at 48-, 56-, 72-, and 80-hour post fertilization. In addition, a comparison between biological models was performed, using results obtained for YFP-C10H-Ras and wild-type YFP-H-Ras for both zebrafish embryos and HEK293T cells. Intragroup variability analysis was carried out between all experimental groups belonging to the same biological model and construct expressed. Initial diffusion coefficients and confinement area sizes were obtained through a confined model fit, by pooling and averaging values of mean square displacements for each individual H-Ras construct, biological model, and time post-fertilization.

In order to check if data were normally distributed, a Shapiro-Wilk statistical test was performed. Significance of the results was performed using a Student’s t-test for comparison of means between two, normally distributed, groups. When multiple groups were compared, a one-way ANOVA was implemented with a Tukey range test for post hoc analysis. Ultimately, the source of potential variability of the data was examined by using a hierarchical linear model to account for nested structure of the data (i.e., different areas within an embryo, different embryos within an experimental day). The values of the *r*^2^ were logarithm-transformed and the *α* logit-transformed to meet the hypotheses of the statistical model.

## AUTHOR CONTRIBUTIONS

Conceptualization: M.J.M.S.; Methodology: R.J.G., B.J., M.A.R., T.S., M.J.M.S.; Software: R.J.G., B.J., P.H, J.N.; Validation: R.J.G., M.J.M.S.; Formal analysis: R.J.G., F.S., P.H.; Investigation: R.J.G., M.A.R, F.S., P.H.; Resources: M.J.M.S.; Data curation: R.J.G., M.J.M.S.; Writing - original draft: R.J.G., M.J.M.S.; Writing - review & editing: R.J.G., J.N., T.S., M.J.M.S.; Supervision: M.J.M.S.; Project administration: M.J.M.S.; Funding acquisition: M.J.M.S.

## REFERENCES

Bobroff, N. (1986). Position measurement with a resolution and noise-limited instrument. Rev. Sci. Instrum. 57, 1152–1157.

Brunsveld, L., Waldmann, H. and Huster, D. (2009). Membrane binding of lipidated Ras peptides and proteins — The structural point of view. Biochim. Biophys. Acta BBA - Biomembr. 1788, 273–288.

Canedo, A. and Rocha, T. L. (2021). Zebrafish (Danio rerio) using as model for genotoxicity and DNA repair assessments: Historical review, current status and trends. Sci. Total Environ. 762, 144084.

Chudakov, D. M., Matz, M. V., Lukyanov, S. and Lukyanov, K. A. (2010). Fluorescent Proteins and Their Applications in Imaging Living Cells and Tissues. Physiol. Rev. 90, 1103–1163.

Dadke, S. and Chernoff, J. (2003). Protein-tyrosine Phosphatase 1B Mediates the Effects of Insulin on the Actin Cytoskeleton in Immortalized Fibroblasts*. J. Biol. Chem. 278, 40607–40611.

Detrich, H. W., III, H. W. D., Westerfield, M. and Zon, L. I. (2011). The Zebrafish: Genetics, Genomics and Informatics. Academic Press.

Eggeling, C., Ringemann, C., Medda, R., Schwarzmann, G., Sandhoff, K., Polyakova, S., Belov, V. N., Hein, B., von Middendorff, C., Schönle, A., et al. (2009). Direct observation of the nanoscale dynamics of membrane lipids in a living cell. Nature 457, 1159–1162.

Garcia, G. R., Noyes, P. D. and Tanguay, R. L. (2016). Advancements in zebrafish applications for 21st century toxicology. Pharmacol. Ther. 161, 11–21.

Garcia-Parajo, M. F., Cambi, A., Torreno-Pina, J. A., Thompson, N. and Jacobson, K. (2014). Nanoclustering as a dominant feature of plasma membrane organization. J. Cell Sci. 127, 4995–5005.

Gheber, L. A. (2018). The Life of a Membrane Protein. Biophys. J. 114, 2762–2763.

Gore, A. V., Pillay, L. M., Galanternik, M. V. and Weinstein, B. M. (2018). The zebrafish: A fintastic model for hematopoietic development and disease. WIREs Dev. Biol. 7, e312.

Groeneweg, F. L., Royen, M. E. van, Fenz, S., Keizer, V. I. P., Geverts, B., Prins, J., Kloet, E. R. de, Houtsmuller, A. B., Schmidt, T. S. and Schaaf, M. J. M. (2014). Quantitation of Glucocorticoid Receptor DNA-Binding Dynamics by Single-Molecule Microscopy and FRAP. PLOS ONE 9, e90532.

Haffter, P., Granato, M., Brand, M., Mullins, M. C., Hammerschmidt, M., Kane, D. A., Odenthal, J., van Eeden, F. J., Jiang, Y. J., Heisenberg, C. P., et al. (1996). The identification of genes with unique and essential functions in the development of the zebrafish, Danio rerio. Development 123, 1–36.

Harms, G. S., Sonnleitner, M., Schütz, G. J., Gruber, H. J. and Schmidt, Th. (1999). Single-Molecule Anisotropy Imaging. Biophys. J. 77, 2864–2870.

Harms, G. S., Cognet, L., Lommerse, P. H. M., Blab, G. A. and Schmidt, T. (2001). Autofluorescent Proteins in Single-Molecule Research: Applications to Live Cell Imaging Microscopy. Biophys. J. 80, 2396–2408.

Hobbs, G. A., Der, C. J. and Rossman, K. L. (2016). RAS isoforms and mutations in cancer at a glance. J. Cell Sci. 129, 1287–1292.

Jacobson, K., Liu, P. and Lagerholm, B. C. (2019). The Lateral Organization and Mobility of Plasma Membrane Components. Cell 177, 806–819.

Kaufman, C. K., Mosimann, C., Fan, Z. P., Yang, S., Thomas, A. J., Ablain, J., Tan, J. L., Fogley, R. D., Rooijen, E. van, Hagedorn, E. J., et al. (2016). A zebrafish melanoma model reveals emergence of neural crest identity during melanoma initiation. Science 351,.

Kawakami, A., Fukazawa, T. and Takeda, H. (2004). Early fin primordia of zebrafish larvae regenerate by a similar growth control mechanism with adult regeneration. Dev. Dyn. 231, 693–699.

Kusumi, A., Sako, Y. and Yamamoto, M. (1993). Confined lateral diffusion of membrane receptors as studied by single particle tracking (nanovid microscopy). Effects of calcium-induced differentiation in cultured epithelial cells. Biophys. J. 65, 2021–2040.

Kusumi, A., Fujiwara, T. K., Chadda, R., Xie, M., Tsunoyama, T. A., Kalay, Z., Kasai, R. S. and Suzuki, K. G. N. (2012). Dynamic Organizing Principles of the Plasma Membrane that Regulate Signal Transduction: Commemorating the Fortieth Anniversary of Singer and Nicolson’s Fluid-Mosaic Model. Annu. Rev. Cell Dev. Biol. 28, 215–250.

Lieschke, G. J. and Currie, P. D. (2007). Animal models of human disease: zebrafish swim into view. Nat. Rev. Genet. 8, 353–367.

Lingwood, D. and Simons, K. (2010). Lipid Rafts As a Membrane-Organizing Principle. Science 327, 46–50.

Lommerse, P. H. M., Blab, G. A., Cognet, L., Harms, G. S., Snaar-Jagalska, B. E., Spaink, H. P. and Schmidt, T. (2004). Single-Molecule Imaging of the H-Ras Membrane-Anchor Reveals Domains in the Cytoplasmic Leaflet of the Cell Membrane. Biophys. J. 86, 609–616.

Lommerse, P. H. M., Snaar-Jagalska, B. E., Spaink, H. P. and Schmidt, T. (2005). Single-molecule diffusion measurements of H-Ras at the plasma membrane of live cells reveal microdomain localization upon activation. J. Cell Sci. 118, 1799–1809.

Lommerse, P. H. M., Vastenhoud, K., Pirinen, N. J., Magee, A. I., Spaink, H. P. and Schmidt, T. (2006). Single-Molecule Diffusion Reveals Similar Mobility for the Lck, H-Ras, and K-Ras Membrane Anchors. Biophys. J. 91, 1090–1097.

Malumbres, M. and Barbacid, M. (2003). RAS oncogenes: the first 30 years. Nat. Rev. Cancer 3, 459–465.

Murakoshi, H., Iino, R., Kobayashi, T., Fujiwara, T., Ohshima, C., Yoshimura, A. and Kusumi, A. (2004). Single-molecule imaging analysis of Ras activation in living cells. Proc. Natl. Acad. Sci. 101, 7317–7322.

Nickels, J. D., Smith, J. C. and Cheng, X. (2015). Lateral organization, bilayer asymmetry, and inter-leaflet coupling of biological membranes. Chem. Phys. Lipids 192, 87–99.

Pezzarossa, A., Zosel, F. and Schmidt, T. (2015). Visualization of HRas Domains in the Plasma Membrane of Fibroblasts. Biophys. J. 108, 1870–1877.

Plowman, S. J., Muncke, C., Parton, R. G. and Hancock, J. F. (2005). H-ras, K-ras, and inner plasma membrane raft proteins operate in nanoclusters with differential dependence on the actin cytoskeleton. Proc. Natl. Acad. Sci. 102, 15500–15505.

Prior, I. A., Muncke, C., Parton, R. G. and Hancock, J. F. (2003). Direct visualization of Ras proteins in spatially distinct cell surface microdomains. J. Cell Biol. 160, 165–170.

Reisser, M., Palmer, A., Popp, A. P., Jahn, C., Weidinger, G. and Gebhardt, J. C. M. (2018). Single-molecule imaging correlates decreasing nuclear volume with increasing TF-chromatin associations during zebrafish development. Nat. Commun. 9, 5218.

Rotblat, B., Prior, I. A., Muncke, C., Parton, R. G., Kloog, Y., Henis, Y. I. and Hancock, J. F. (2004). Three Separable Domains Regulate GTP-Dependent Association of H-ras with the Plasma Membrane. Mol. Cell. Biol. 24, 6799–6810.

Schaaf, M. J. M., Koopmans, W. J. A., Meckel, T., van Noort, J., Snaar-Jagalska, B. E., Schmidt, T. S. and Spaink, H. P. (2009). Single-Molecule Microscopy Reveals Membrane Microdomain Organization of Cells in a Living Vertebrate. Biophys. J. 97, 1206–1214.

Schütz, G. J., Schindler, H. and Schmidt, T. (1997). Single-molecule microscopy on model membranes reveals anomalous diffusion. Biophys. J. 73, 1073–1080.

Seefeldt, B., Kasper, R., Seidel, T., Tinnefeld, P., Dietz, K.-J., Heilemann, M. and Sauer, M. (2008). Fluorescent proteins for single-molecule fluorescence applications. J. Biophotonics 1, 74–82.

Semrau, S. and Schmidt, T. (2007). Particle Image Correlation Spectroscopy (PICS): Retrieving Nanometer-Scale Correlations from High-Density Single-Molecule Position Data. Biophys. J. 92, 613–621.

Simons, K. and Gerl, M. J. (2010). Revitalizing membrane rafts: new tools and insights. Nat. Rev. Mol. Cell Biol. 11, 688–699.

Singer, S. J. and Nicolson, G. L. (1972). The Fluid Mosaic Model of the Structure of Cell Membranes. Science 175, 720–731.

Willumsen, B. M., Christensen, A., Hubbert, N. L., Papageorge, A. G. and Lowy, D. R. (1984). The p21 ras C-terminus is required for transformation and membrane association. Nature 310, 583–586.

